# First Detection of Pathogenic *Escherichia coli* Isolates Associated With Donkey Foals’ Diarrhea in Northern China

**DOI:** 10.1101/2020.04.06.027458

**Authors:** Liu Wen-qiang, Xia Nan, Zhang Jing-wen, Wang Ren-hu, Jiang Gui-miao

**Affiliations:** College of Agriculture, Liaocheng University, Liaocheng, Shandong, 252000, China; Donge ejiao co.,Ltd,Liaocheng, Shandong, 252000, China

**Keywords:** donkey, *E.coli*, identification, influence factor, genome

## Abstract

**Objective:** The aim of this study was to identify the biological features, influence factor and Genome-wide properties of pathogenic donkey *Escherichia coli* (DEC) isolates associated with severe diarrhea in Northern China.

**Methods:** The isolation and identification of DEC isolates were carried out by the conventional isolation、automatic biochemical analysis system、serotype identification、16S rRNA test、animal challenge and antibiotics sensitivity examination. The main virulence factors were identified by PCR. The complete genomic re-sequence and frame-sequence were analyzed.

**Results:** 216 strains of DEC were isolated from diarrhea samples, conforming to the bacterial morphology and biochemical characteristics of *E*.*coli*. The average size of the pure culture was 329.4 nm×223.5 nm. Agglutination test showed that O78 (117/179, 65.4%) was the dominant serotype and ETEC(130/216, 60.1%) was the dominant pathogenic type. Noticeable pathogenic were observed in 9 of 10 (90%) randomly selected DEC isolates caused the death of test mice (100%, 5/5) within 6h∼48h, 1 of 10 (10%) isolates caused the death of test mice (40%, 2/5) within 72h. Our data confirmed that DEC plays an etiology role in dirarrea/death case of donkey foal. Antibiotics sensitivity test showed significant susceptibility to DEC isolates were concentrated in Nor、EFT、ENR、CIP and AMK,while the isolates with severe antibiotic resistance was AM、TE、APR、FFC、RL and CN. Multi-drug resistance was also observed. A total of 15 virulence gene fragments were determined from DEC(n=30) including OMPA (73%), safD (77%), traTa (73%), STa(67%), EAST1 (67%), astA (63%), kspII (60%), irp2 (73%), iucD (57%), eaeA (57%), VAT (47%), iss (33%), cva (27%), ETT2 (73%) and K88 (60%) respectively. More than 10 virulence genes from 9 of 30(30%) DEC strains were detected, while 6 of 30(20%) DEC strains detected 6 virulence factors. phylogenetic evolutionary tree of 16S rRNA gene from different isolates shows some variability. The original data volume obtained from the genome re-sequencing of DEC La18 was 2.55G and Genome framework sequencing was carried out to demonstrate the predicted functions and evolutionary direction and genetic relationships with other animal *E*.*coli*.

**Conclusions:** These findings provide firstly fundamental data that might be useful in further study of the role of DEC and provide a new understanding of the hazards of traditional *colibacillosis* due to the appear of new production models.

## 1. Introductions

It is well known that *E. coli, salmonella, clostridium* and *rotavirus* are the main pathogenic factors causing diarrhea in infants、poultry and young animals^[1]^. Evidence suggests that specific serotype of *E*.*coli* is among the most important factors for enteric disease and sepsis^[2-4]^. The resulting problems of livestock and poultry deaths, antibiotic residues and the spread of drug resistance become a growing public health concern worldwide. There are relatively few historical studies in the area of Equine *colibacillosis. E*.*coli* infection in Equine family has mainly sporadic and serious morbidity was not noticed under natural farming previously^[5]^. Treatment for individuals of the colt is effective and practicable. However, more than 200 donkey farms had been established around Liaocheng city with a stock of nearly 100,000 and replicated in other provinces in northern china quickly since 2015 (China Statistical Yearbook 2015-2018). The issue has grown in importance in light of recent the rapid new built donkey farms. These rapid changes are having a serious effect on the transmission、propagation and epidemic of pathogenic *E*.*coli*^[6]^. Recently clinical observation suggested that diarrhea was the most frequently occurred disease with average morbidity of newborn colts (under 3 months age) was 23.1% (1099/4755), about 15% of the colts died within 48 to 72 hours of onset^[6,7]^. In light of recent events in serious economic losses, it is becoming extremely difficult to ignore the existence of pathogenic *E*.*coli* in donkey herding. the veteterinary should give great attention to the prevalence of *colibacillosis*. The purpose of this thesis is to document the etiological studies of DEC strains and contribute to this growing area of research by exploring the features of *colibacillosis* in donkey farms.

## 2. Materials and methods

### 2.1 Recording of cases

All the samples are collected in 19 intensive Dezhou donkey farms (the stock is between 300 and 1000) newly built since Jan 2016 located in northern China. Criteria for selecting the subjects were as follows: diarrhoea of foals less than 3 months old^[7, 8]^. Totally, 363 cases and death (n=108) due to severe diarrhea were concentrated within among them were recorded. Samples from the sicked or death foals were collected include liver 、diseased intestinal contents and anal swab according to the standard method^[9]^. Briefly, the samples were collected with sterile cotton swabs that were placed into semi-solid Amie’s transport medium and transferred to the Research Institute of Donkey Breeding and Feeding of Liaocheng University.

### 2.2 Culture and identification of bacteria strains

Pathogenic bacteria from the sample were cultured by inoculating the bacteria in 5% calf serum nutrient agar medium, MacConkey medium(Invitrogen Thermo Fisher Scientific-CN) with aseptic techniques according to the standard schedule^[10]^. Cultures were aerobically incubated at 37°C for 24h, and the degree of bacterial growth was assessed semi-quantitatively. 3∼4 clones from each plate were selected and adjust the concentration of single colony bacteria to 0.5 McKlisterian turbidity. All the clones were analyzed using the micro-biochemical reaction manufacturer (Shandong Xinke Biotech) to determine the species of the *E.coli* strains. 216 strains of DEC were determined in the sicked samples and were stored in LB liquid media with equal volums sterilized glycerin at -80°C totally. ATCC25922 as the reference strain from China Institute of Veterinary Drugs Control(CIVDC)^[10, 11]^.

### 2.3 Ethics statement

Mouse studies were performed accordance with the recommendation in the Guide for the Care and Use of Laboratory Animals of the Ministry of Science and Technology of PRC. The animal carcasses are sent to the nearest harmless treatment plant after the samples are collected. Animal research approval was obtained from the Animal Welfare Committee of Liaocheng University.

### 2.4 Electron microscopic morphology of typical E.coli

DEC La18 strains with typical biological characteristics and good pathogenicity was registered for preservation in CGMCC (No.14058) and selected for the following analysis of morphological characteristics. Briefly^[10,11]^, 20μl DEC La18 strains bacterial suspension of 0.5 MCF prepared was added to the copper net with supporting film. After standing for 15 min, the excess liquid was extracted and slightly dried; then, a drop of 2% phosphotungstic acid negative dye solution was added and standing for 10 min and dried naturally. Last, samples were observed by transmission electron microscope and photographed.

### 2.5 Identification by 16s rRNA sequence

Genomic DNA samples of 30 DEC isolates randomly selected from 216 isolates were separated with SK8255 DNA extraction kit. Universal primers for *E.coli* identification of 27F (5’ -agagtttgatcctggctcag-3’) and 1492R (5’ -tacggctaccttgttacgtt-3‘) were amplified and for PCR process^[12, 13]^. Then, 16s rRNA PCR products about 1500 bp was sequenced (Sangon Biotech) and re-confirmed with the identification results. The species of DEC isolates were determined with screening in ribosomal database http://rdp.cme.msu.edu/index.jsp. The representative nucleotide sequence (n=10) were registered in GenBank.

### 2.6 Mice challenge experiments and microbiology detection

isolate strains confirmed in *2.5* was selected randomly for mice challenge to determine the pathogenicity. Follow the following procedure each time^[14]^: five Four-week-old BALB/C mice (average 25g body weight) were challenged with intraperitoneal injection by 5×10^8^ CFU in a volume of 0.2mL. The control group was injected with the same amount of sterile LB culture medium. All animals are raised at room temperature. Clinical symptoms include depressed of spirit 、temperature changes、loss of appetite/waste、diarrhea and the mortality of mice were recorded statistically from 6h, 12h, 18h, 24h, 36h, 48h, 72h after challenge. The livers of mice were autopsied, and surviving mice euthanized under inhalation anesthesia after 72h, The homologous bacteria were identified again by microscopy and micro-biochemical reactions according to the methods described above.

### 2.7 Antimicrobiology susceptibility testing

drug-sensitive paper including Ampicillin (AM), Norfloxacin (Nor), Ceftiofur (EFT), Ciprofloxacin (CIP), Sulfamethoxazole (RL), Gentamicin (CN), Enrofloxaci (ENR), Tetracycline (TE), Kanamycin (AMK), apramycin (APR) and florfenicol (FFC) were performed to the antibiotic sensitivity test for the identified strains (n=216) accordance with the standard procedure of Kirby-Bauer method recommended by CLSI through the measure of inhibition zone diameters for each antimicrobial tested^[14]^.

### 2.8 Determination of serotype and pathotype

Mono-factor serum (specific for ETEC、STEC、EPEC、EAEC and EIEC) was used as aggregative adherence assays. Antigen samples prepared (n=216) were respectively tested with somatic O antigen (O1-O185) serum (CIVDC) using standard agglutination methods to determine the distribution of DEC serotypes^[15, 16]^.

### 2.9 Classification of virulence genes using PCR assays

A total of 37 primers for the major virulence genes were pre-designed for PCR. Tab1 showed detailed information on the primers used for screening for virulence genes^[2, 5, 7, 9, 11, 17, 31]^. 30 isolate strains same as from *2.5* for PCR assay. Briefly, Monoclone bacteria were grown on Tripticase Soy Agar (Difco) at 37°Cfor 24h. DNA was extracted by suspending three bacterial colonies in 200 µl of sterile water that was boiled for 10 min and centrifuged at 10,000 g for 5 min. The supernatant was used as the template in the PCR assays. The presence of virulence genes was examined using PCR techniques described previously. Isolates positive for *eae*+ were typed according to previously described criteria^[17, 20]^.

### 2.10 Analysis of genomic sequence

Genomic DNA was extracted from DEC La18 strains for analysis of gonomic sequence by re-sequencing and frame sequencing (Sangon Tech)^[13, 21, 22]^. The genomic data were registered in Genbank and analyzed comparing with the reference genome sequences of equine *E.coli* strain (3.2608 Contig0022) and bovine *E.coli* O157:H7 strain (FRIK2455) respectively. Based on Illumina HiSeq platform, the whole genome framework of DEC La18 was sequenced, genome splicing and annotation circle mapping were performed, and the functions of virulence genes、drug resistance genes and host-pathogen interaction were annotated. BLAST was used to compare the gene protein sequences obtained with the pathogen-host interaction database, so as to obtain the interaction information between the gene and its corresponding pathogen-host.

## 3. RESULTS

### 3.1 isolation and identification of DEC

216 strains of DEC were determined by microbiological methods in the sicked samples totally. 94 of 216 (43.5%) strains were separated from 108 cases of death after diarrhoea, while the remaining 122 (56.5%) were isolated from 255 cases of non-death after diarrhoea. Approximately, the isolation rate was 59.5% from 363 cases of diarrhea and death. Other suspected bacteria and virus are not presented in this paper. Pink bacterial colonies with smooth and neat edges in McConkey medium was cultured. Black colonies with metallic luster, moist surface and green edge in Eosin Methylene Mlue Agar medium was observed (fig1). The isolates were consistent with the micro-biochemical reaction *E.coli* results of 99% system confidence. The average size of the pure culture of DEC La18 strain was calculated about 629.4nm×223.5nm. Transmission electron microscopy observed the presence of bacteria individually or in pairs (fig2-1), fimbriea of bacteria were also remarkable observed (fig2-2).

**Fig 1.**
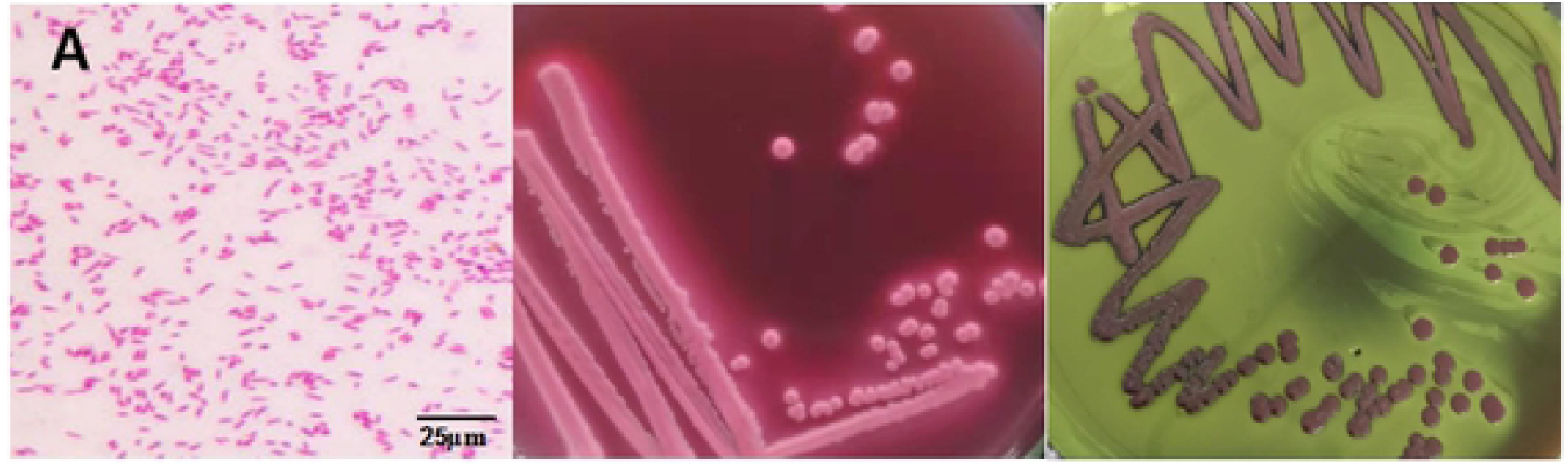
G-(10×40), pink bacterial colonies in McConkey medium and black colonies in Eosin Melhylene Mlue Agar medium.

**Figure 2-1.**
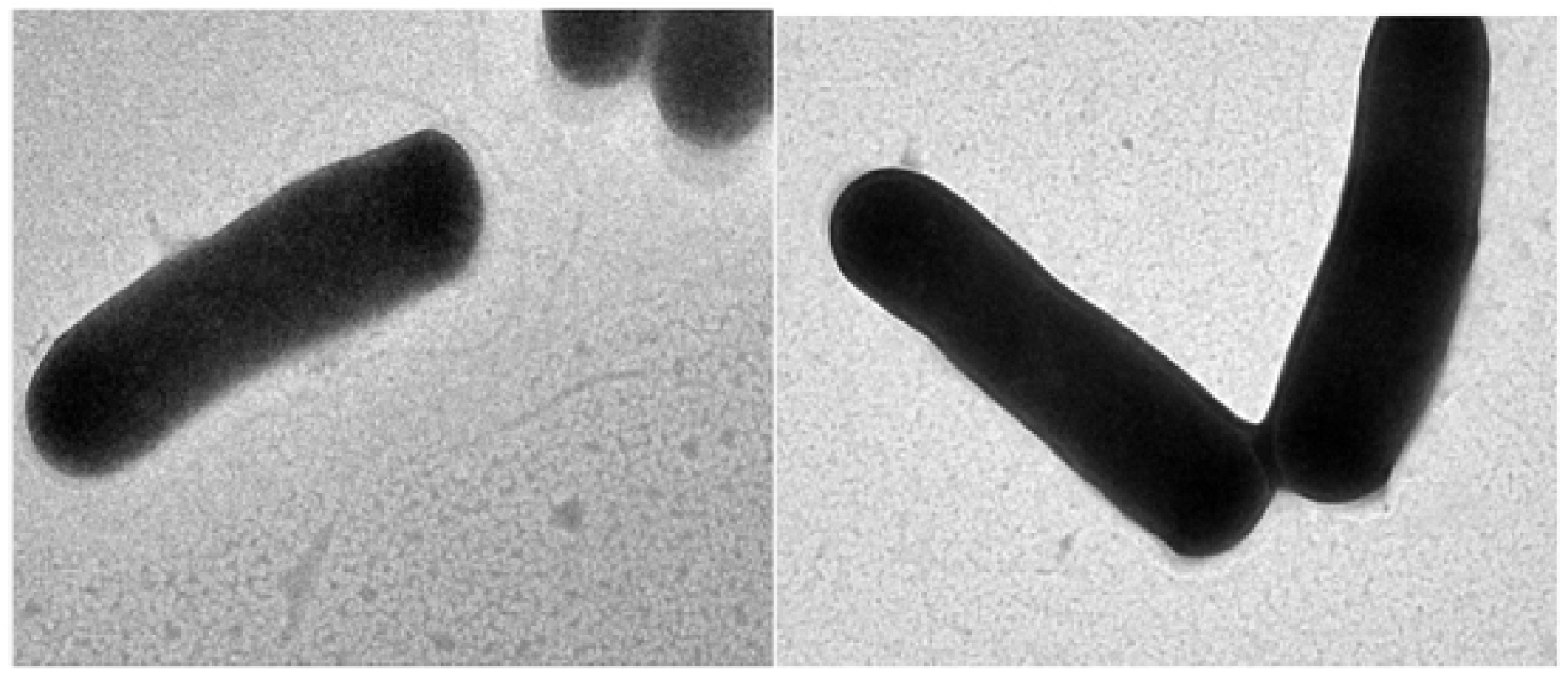
Morphology of isolates under transmission electron microscopy(10000×)

**Fig 2-2.**
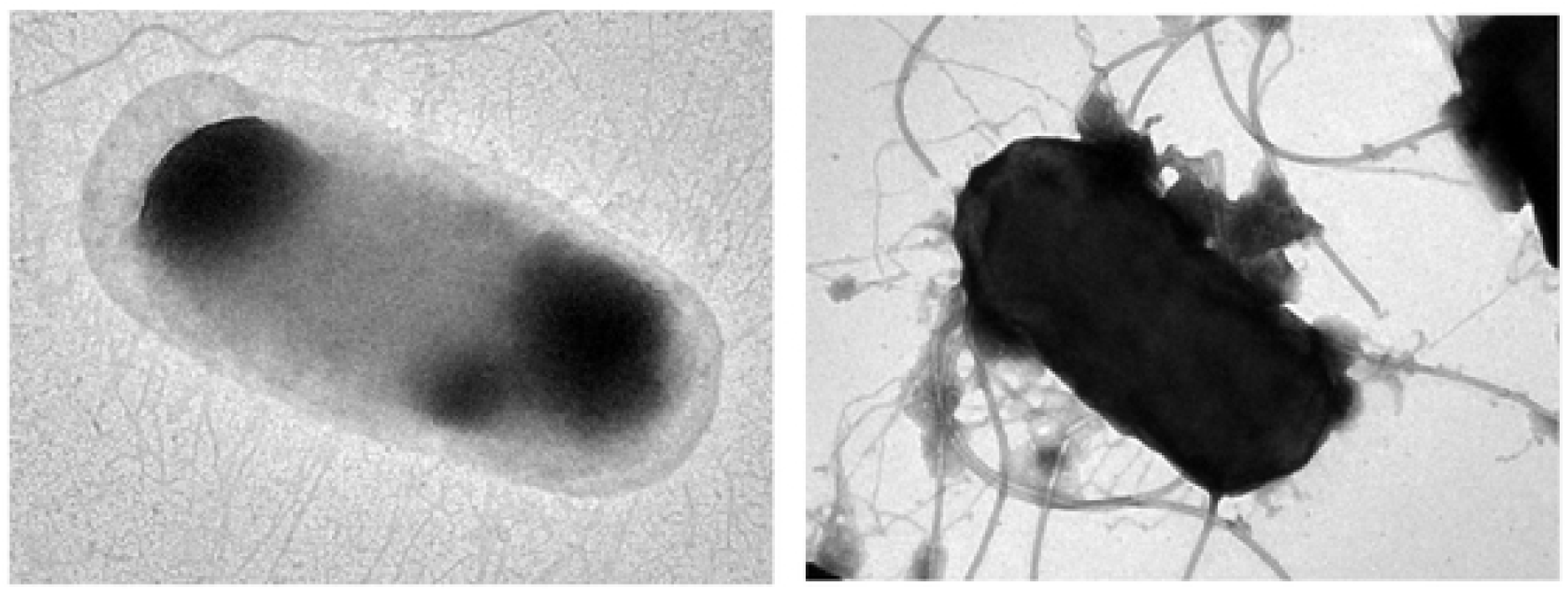
Slained pili observation in transmission electron microscopy(30000×)

### 3.2 Analysisi and phylogenetic tree of DEC 16s rRNA

The 16s homology of all ass *E.coli* was more than 98.9%, but phylogenetic evolutionary tree of 16S rRNA gene from different isolates show some variability. MF104548 and MN083305 appear to form a similar branch with homologue from cattle and sheep, while MF104543 was located a similar branch with homologue from human and horse (fig3).

**Fig 3.**
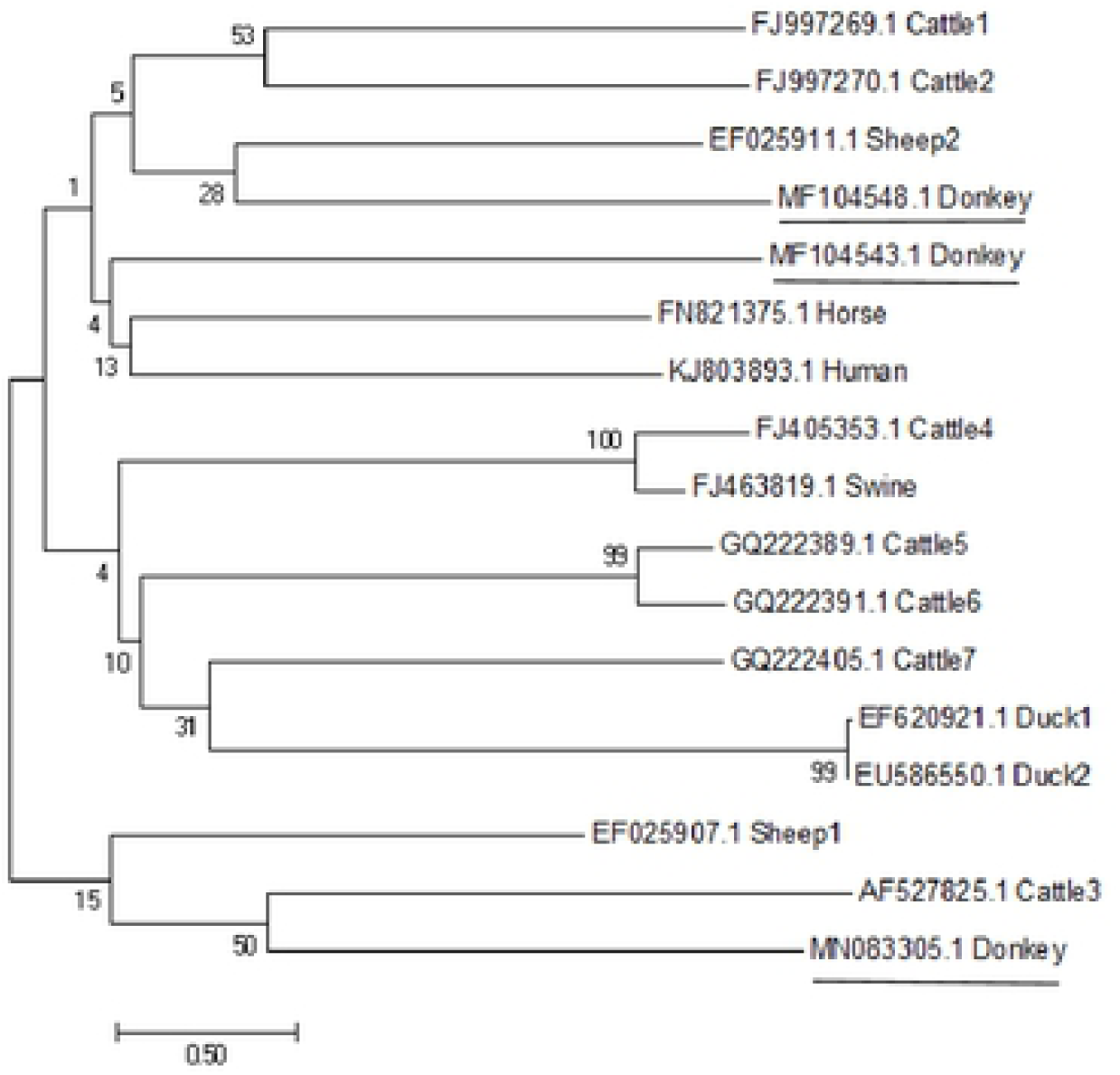
Genetic evolutionary tree analysis of 16S rRNA gene from different source

### 3.3 Pathogenicity of several representative isolates

Clinical symptoms include elevated body temperature (fig4), lethargy, loss of appetite, shortness of breath and diarrhea was observed as early start as 6h in two groups challenged by La18 and Lb37. All the challenged animals developed the obvious symptoms after 18 h. The first mice died after 6h and others 4 animals died within 12h in DEC La18 challenge group. Animal deaths in other 8 groups were relatively concentrated in 12 h∼24 h. The latest death mice was recorded at 48h after challenged by DEC Lb9. Although typical clinical symptoms were observed in DEC La44 challenge mice, the death rate was the lowest (2/5). No abnormality was found in the control group animals. The typical histopathological changes of the mice after autopsy are shown in fig5. This results indicate that the isolated bacteria had significant pathogenicity.

**Fig 4.**
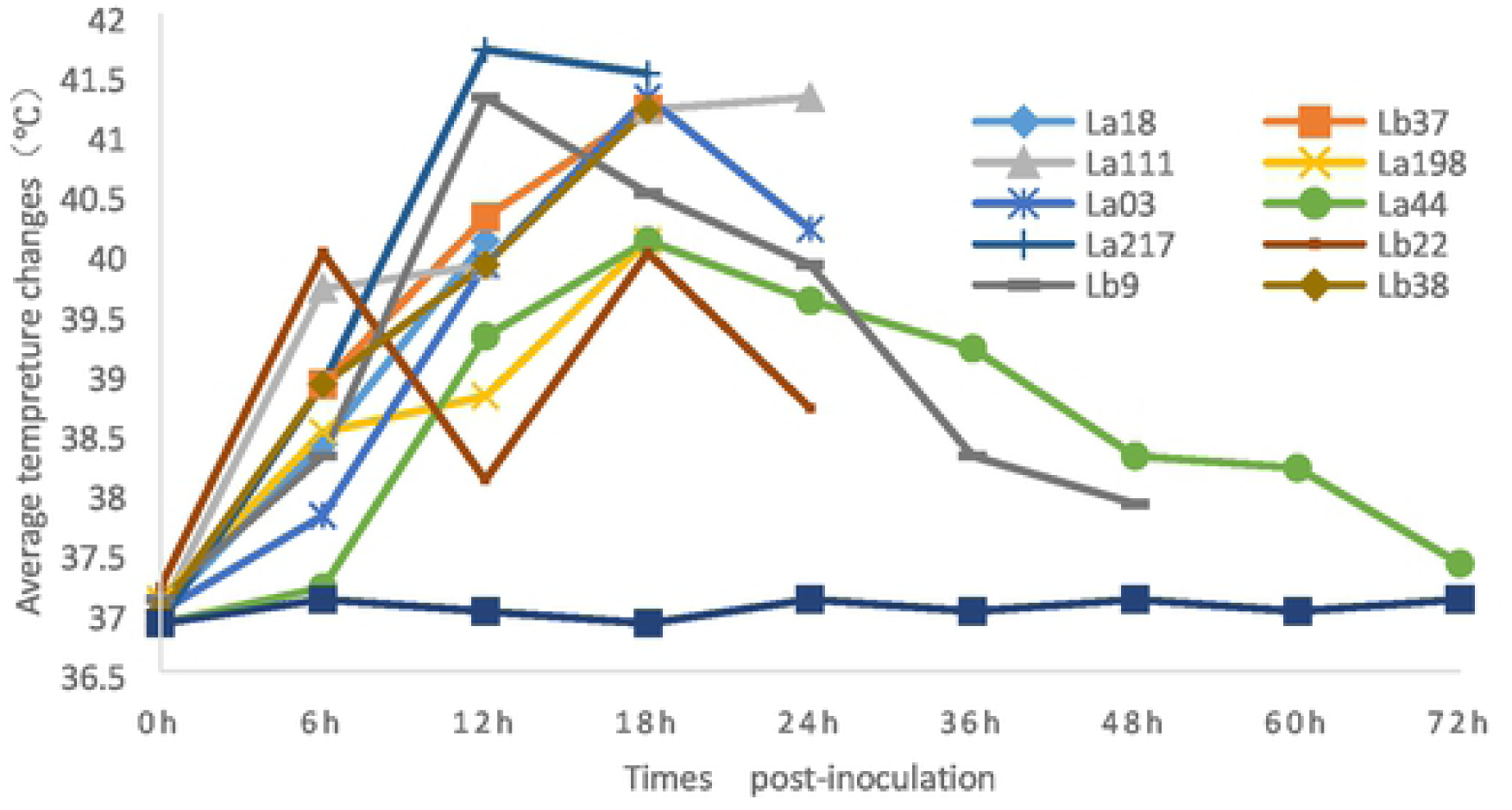
average temprature changes or post-inoculation until all the animals in the groups

**Fig 5.**
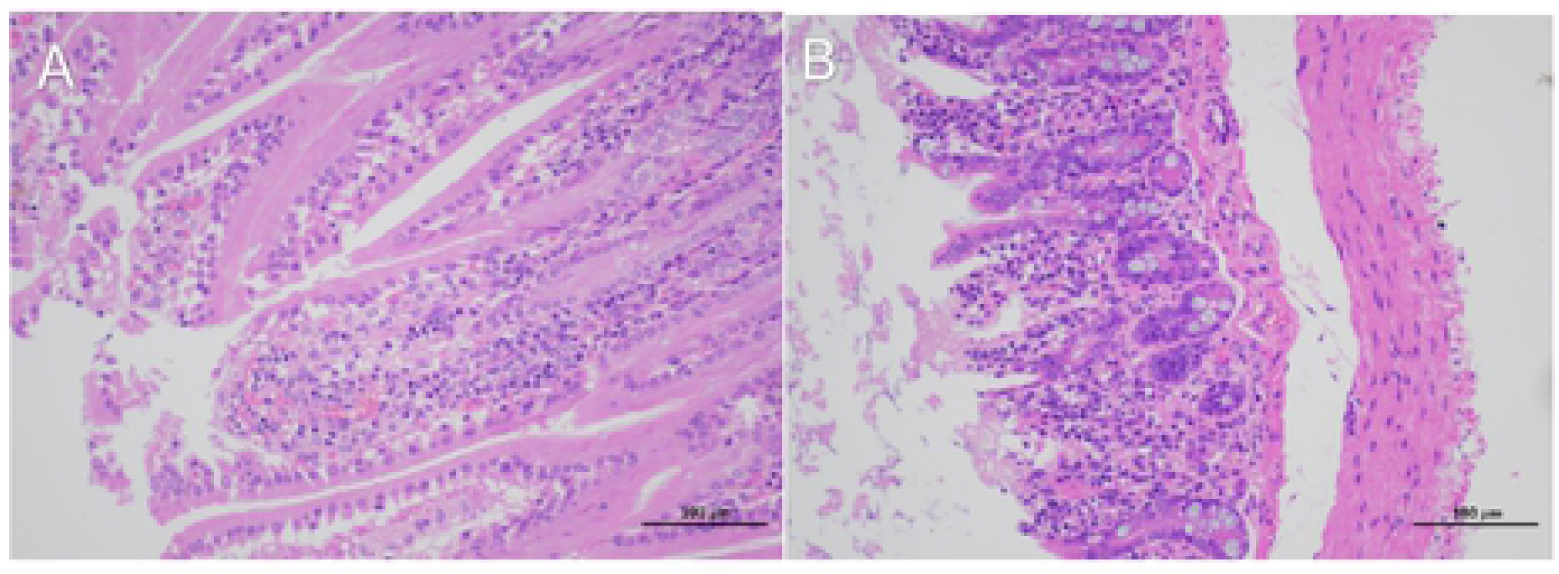
Pathological section or mouse intestinal tract shows that Intestinal mucosal epithelium was seriously detached, congestion of the duodenum (A) and The muscularis and outer membrane or ileum were seriously damaged (B)

### 3.4 Antibiotic sensitivity test results

216 DEC isolates showed different antibiotics resistance rates to 11 commonly used antibiotics in animals of China are as follows: AM (88.4%)、Nor (47.5%)、EFT (36.2%)、CIP (45.4%)、RL (79.5%)、CN (93.1%)、ENR (51.9%)、TE (83.6%)、AMK (51.9%) 、APR (67.1%) 、FFC (93.1%). Antibiotics with high susceptibility to all isolates were concentrated in Nor、EFT、ENR、CIP and AMK,while the isolates with severe antibiotic resistance was AM、TE、APR、FFC、RL and CN (tab 2). 77 of 216(35.6%) DEC strains showed resistance to five or more antimicrobial agents among them.

### 3.5 Serotype and pathotype

The serotype of 179 isolates (82.9%) from the selected 216 isolates strains selected was identified as the O78 (117, 65.4%) 、O101 (37, 20.7%) and O8 (25, 13.9%) respectively. Profile of ETEC was found in 86 of 94 (91.5%) with diarrhea/death, and 44 of 122 (36.1%) with diarrhea without death, the overall ratio is 60.1%(tab2). Profile of EPEC was found in 8 of 94 (8.5%) with diarrhea/death, and 35 of 122 (28.7%) with diarrhea without death, the overall ratio is 20.4%. No results were obtained from 42 samples (19.4%).

### 3.6 Classification and distribution of major virulence factors

PCR results showed that a total of 15 virulence gene fragments were obtained from 30 strains of DEC. The positive rate of virulence factors of OMPA (73%), safD (77%), traTa (73%), STa (67%), EAST1 (67%), astA (63%), kspII (60%), irp2 (73%), iucD (57%), eaeA (57%), VAT (47%), iss (33%), cva (27%), ETT2 (73%) and K88 (60%) were determined respectively. The genes of other virulence factors were not detected. More than 10 virulence genes from 9 of 30 (30%) DEC strains were detected, the least of them, 6 of 30 (20%) strains of DEC all detected 6 virulence factors, while one strain named La18 carries up to 13 virulence factors (OMPA, safD, ETT2, traTa, sta, EASTI, astA, KSPII, irp2, iucD, VAT, TSH, iss) especially (tab3).

### 3.7 Whole-genome analyses of DEC

3.7.1 The original data volume obtained from the genome resequencing of DE La18 was 2.55G (genebank registered number SRP148724) with 51% G+C base content and average sequencing depth 229.3 of all bases. Whole genome includ 35586 SNP and 4157 InDel, comparing with the reference genome sequences of equine *E.coli* strain (3.2608 Contig0022) and bovine *E.coli* O157:H7 strain (FRIK2455) respectively. Compared with equine and bovine *E.coli*, the homology of DEC genome is 98.29% and 96.67%, respectively. Mutations were most concentrated in the NZ-AEZS000010.1 chromosome. *Downsteam-gene-variant* 、*Trompe-variant* and *Trompe-gene-variant* of the 13 mutation types had the highest mutation counts, reaching 175,222, 27,751, and 178,117 respectively, distributing in the 3 corresponding mutations in the highest number of genomic regions including *Downstream、Exon* and *Upstream*.

3.7.2 Genome framework sequencing (Genbank registered number PRJNA587570) was predicted to contain 4,544 genes, of which 3,247 were more than 500bp in length, and 1,626 were more than 1,000 bps in length, with an average gene length of 913. The framework diagram of the COG classification of the genome are shown in fig6. There are 4259 transcripts with annotation information in NR database. The predicted proteins were annotated to 343 (7.9%,SetA) and 383 (8.8%,SetB) virulence factor proteins in the VFDB database, respectively. 108 predicted antibiotics resistance proteins of 4320 (2.5%) was identified. In 212 of 4320 (4.91%) PHI information, participate in “reduced virulence process” gene (n=93) 、“unaffected pathogenicity” process gene (n=80)、effector gene(n=25)、”loss of pathogenicity”gene(n=23)、”increased virulence” genes(n=14) 、“mixed outcome” process genes (n=11) 、“lethal” genes (n=6) and “Resistance to chemical” process 1 gene were listed.

**Fig 6.**
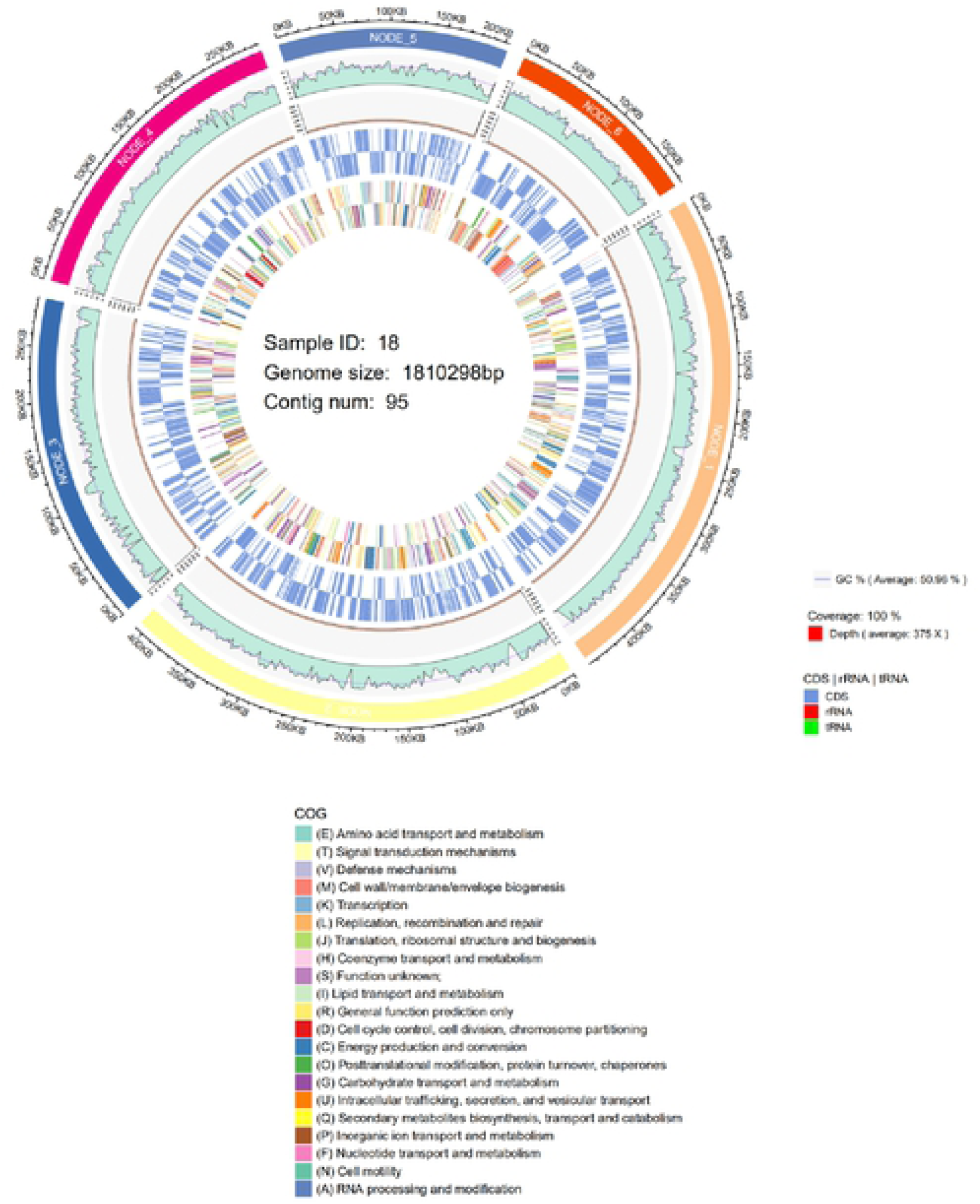
The framework diagram of the genome of DEC La18 strain

## 4. DISCUSSION

Equine sourced *E. coli* is generally used as an important target bacteria that focus on drug resistance research and environmental observation^[23]^. Very little was found in the literature on the question of *Colibacillosis* with serious infections or epidemics in the equine family. In the state of natural culture as a driving vehicle, the serious incidence of donkey *Colibacillosis* is rare but serious occurs naturally in large-scale farms. Therefore, the initial goal of the objective is to identify the etiological features of DEC isolates associated with the donkey *Colibacillosis* characterized by serious diarrhea and high mortality of foals in recent years. Based on the etiology of pathogenic *E.coli*, 216 isolates were identified as associated with the samples from sicked donkeys in northern China. Therefore, based on the definite etiologic diagnosis including bacterial morphology, biochemical reaction, animal challenge test, determination of 16srRNA, the following conclusion can be confirmed that DEC with high pathogenicity *was* the single agent of the cases of diarrhea and death of donkey colts in large-scale donkey farms.

Before this study, evidence of donkey epidemic disease was not concerned except as an adjunct study to equine zoonotic diseases. The contribution of this study has been to confirm the occurrence pattern of zoonotic diseases: the aggregation of animals makes the original sporadic individual infection become epidemic case quickly, as previously reported^[7,18]^ and our study.

There have been many reports on the identification of animal and enterobacteria serotypes, but the distribution of the O antigen is relatively stable in certain regions^[16]^. O1, O2 and O78 are the relatively dominant serotypes in chicken^[24]^. 203 (61.70%) of the 329 species were *E.coli* isolated from fecal samples in Nigeria, of which 76 were serotype O157^[25]^. O78 was the dominant DEC serotype in our study, which was significantly different and simpler from the results from cattle, equine and human sources^[5, 9, 16]^. The serotype of *E.coli* O-antigen in foal diarrhea is related to pathogenicity, so it can be helpful for epidemiological investigation and provide a good basis for the preparation of effective vaccines. O157: H7 serotype was not isolated suggesting that no significant risk to food safety although the source of donkey sourced *E.coli* was relatively complex. Although there are a few different samples, we also determined that *K88+*ETEC was the major pathogenic type in DEC. This also accords with our earlier observations^[7]^, which match those previous consensus especially from human beings、swine and cattle diarrheal disease.

Another important finding was that DEC isolates in this study indicate severe antibiotics resistance and high pathogenicity. The isolates were sensitive to antibiotics such as Nor、EFT、ENR、CIP and AMK, while display severe resistant to AM、TE、APR、FFC、RL and CN. This is useful to guide clinical treatment. Through whole-genome sequencing, it was found that the bacteria carried a variety of antibiotic resistance genes, and the expulsive pump system may be the main factor for the mechanism of drug-resistance, consisting with previously report^[25, 27]^. The expulsive pump and its regulatory genes include acrA, acrD, acrE, acrR, acrS, cpxR, emrB, emrR, evgA, h-ns, marA, mdtE, mdtF, mdtG, mdtL, mdtN, mdtO, mdtP, soxR, soxS, tolC. These 20 factors mediate multi-drug resistance of clinical importance and often spread resistance by horizontal transfer of mobile factors between strains of different genera. Rifampicin, Fluoroquinolones, Fosfomycin, Neomycin, Aminoglycosides and other AR genes were also carried in the sequence. Drug-resistant genes also produce antibiotic resistance in other ways, such as participating in bacterial structure construction (bacA, ompF, PmrB, PmrC, PmrF, etc) and participating in various protein or activity processes (arnA, APH (6) -id, MFD).

On the whole, the virulence factor positive rate carried by DEC is different from other reports^[26]^, but higher than the general virulence factor positive rate such as from *E.coli* isolated from animal feces^[27]^. Fifteen virulence factor genes were detected from 30 DEC strains contain the adhesive fimbriae、virulence island、Enterotoxin、antiserum survival factor and secretory system. The high positive rates of adhesive fimbriae K88 and sfaD relating to the invasion and colonization of DEC and play an important colonization role in the pathogenesis of infection. These two virulence factors are also commonly present in diarrheal *E.coli* in piglets. ETT2、eaeA and irp2 are highly detected in weaned pig and cow with *E.coli* infection as a marker gene to judge the presence of HPI in pathogenic bacteria^[28]^. ETT2 virulence island (III secretion system) as the main virulence gene cluster was first found in *E.coli* O157: H7, located near the loci tRNA glyU in the bacterial chromosome. irp2 virulence factor as an iron regulatory gene in highly pathogenic virulence island (HPI), only exists in virulent pathogenic strains and is closely related to virulence and evolution. Enterotoxin gene including SafD、astA、sta and EAST detected in DEC, which is an important pathogenic factor causing abdominal fouling disease in donkeys^[29]^. The high detection rate of above 7 virulence factors indicating that the isolates in our study had significant pathogenicity. This finding was consistent with the results of the animal challenge test. Several important antiserum survival factor including OMP、AtraT、aiss and cva effector genes were detected in DEC^[30, 31]^, which suggested that the existence of these virulence factors is conducive to the survival of bacteria, breaking through the protection and transmission among the donkey farms. VAT was first discovered and reported by Parreira^[18, 27]^, and it exists on the virulence island and encodes 148.3 kDa autonomous transporter (T5SS). VAT is an important part of APEC pathogenicity and has the cytotoxic activity of cell vacuolation. The discovery of VAT distribution in DEC isolates indicates that it also plays an important role in EHTC, which is worth further study.

At the genome-wide level, NCBI genome database has registered more than 170 *E. coli* genome sequence information about strains from poultry、pig、human being. By comparing with the genome, the mutation location, mutation type, mutation gene and chromosome with the severe mutation were found, which provided data support for further study on pathogenetic DEC. Our study helped to enrich the data of *E. coli* in animals, particularly equines. Frame sequencing of genome demonstrated that the matching rates of drug-resistant genes and strains *E. coli* STR.K-12substr MG1655 and *E. coli* STR.K-12substr W3110 were higher (53 and 36, respectively). It indicates that DEC enriched the characteristics of transmissibility 、wide spectrum of drug resistance、strong pathogenicity. Together, these factors contributed to the occurrence of foal *Colibacillosis* characterized by severe diarrhea and high mortality. The information of relevant genes、regulatory elements and predictive proteins obtained by DEC sequencing is helpful for the further study and application of the biological characteristics of the bacteria.

The phylogenetic tree of the 16S rRNA gene from different DEC isolates shows some variability, indicating the complexity of the origins whether it’s from other animals or other areas. These relationships may partly be explained the facts that the donkey mass movement from the northeast and Inner Mongolia to Shandong province in recent years. Nevertheless, such speculations may need more data to back them up.

The main goal of the current study was to determine the biological features of DEC associated with the direct pathogen of diarrhea in newborn donkey foal. The insights gained from this study may be of assistance to attaches great importance to *Colibacillosis* in large-scale donkey farms. Since the study was limited to DEC, it was not possible to isolate other pathogens such as *rotavirus* or *salmonella*. Further research should be carried out to establish effective environmental control methods to reduce the risk of transmission of pathogenic *E.coli*.

## ACKNOWLEDGEMENTS

We are grateful to the owners and veterinarians of the examined donkeys who participated in this research.

## SOURCE OF FUNDS

Donkey Innovation team Project of Modern Agricultural Product Technology system in Shandong Province (SDAIT-27)、Scientific research project of Liaocheng University,318011701

